# Dynamics as a cause for the nanoscale organization of the genome

**DOI:** 10.1101/2020.02.24.963470

**Authors:** R. Barth, G. Fourel, H. A. Shaban

## Abstract

Chromatin ‘blobs’ were recently identified by live super-resolution imaging as pervasive, but transient and dynamic structural entities consisting of a few associating nucleosomes. The origin and functional implications of these blobs are, however, unknown. Following these findings, we explore whether causal relationships exist between parameters characterizing the chromatin blob dynamics and structure, by adapting a framework for spatio-temporal Granger-causality inference. Our analysis reveals that chromatin dynamics is a key determinant of both blob area and local density. However, such causality can only be demonstrated in small areas (10 – 20%) of the nucleus, highlighting that chromatin dynamics and structure at the nanoscale is dominated by stochasticity. Pixels for which the inter-blob distance can be effectively demonstrated to depend on chromatin dynamics appears as clump in the nucleus, and display both a higher blob density and higher local dynamics as compared with the rest of the nucleus. Furthermore, we show that the theory of active semiflexible polymers can be invoked to provide potential mechanisms leading to the organization of chromatin into blobs. Based on active motion-inducing effectors, this framework qualitatively recapitulates experimental observations and predicts that chromatin blobs might be formed stochastically by a collapse of local polymer segments consisting of a few nucleosomes. Our results represent a first step towards elucidating the mechanisms that govern the dynamic and stochastic organization of chromatin in a cell nucleus.

## Introduction

The eukaryotic genome is hierarchically structured from the level of nucleosomes over chromatin loops, topologically associated domains (TADs) and phase-separated A/B compartments up to chromosome territories ^1^. These structural elements are not static but dynamics entities ^2^, such that an appreciable heterogeneity between cells ^3,4^ and in dynamics over time ^2,5^ exist. Such dynamics is borne out from a host of players interacting with the chromatin fiber, including enzymes that typically use ATP for their function associated with a motion component (polymerases ^6^, chromatin remodelers, topoisomerases, helicases, cohesins and condensins ^7–9^, …) as well as mere binders, such as HMGB proteins^10^. We will refer to these proteins as “Active Effectors” in this article. Accordingly, dynamics of the chromatin fiber are altered locally^11,12^, but also globally ^13–18^, in response to nuclear processes such as transcription. Both dynamics and conformational flexibility are key to allow for long-range communication within a fiber, as involved in innumerable genomics processes and in particular in gene activation by transcriptional enhancers.

Given the observed relationships between structural reorganization of the genome^19,20^, nuclear functions and chromatin dynamics, a lasting question in genome biology remains if, and if yes how, chromatin dynamics has an effect on genome organization in nuclear space. To tackle this question, we recently introduced Deep-PALM, a live-cell super-resolution approach able to achieve sub-diffraction spatial resolution and 360 ms temporal resolution to image chromatin *in vivo* ^21^. Using Deep-PALM, chromatin was shown to assemble in nanometer-sized ‘blobs’ (see Figure 1A). These chromatin blobs manifest as accumulation of fluorescence signal from the fluorescently labeled core histone H2B over the background in super-resolution images (Figure 1A). A few (< 30, ref. ^21,22^) nucleosomes likely associate transiently within the time scale of about 1 second, but the functional implications of their existence is not explored. The ability to obtain time-resolved super-resolution images of chromatin allowed to integrate an Optical Flow approach ^14,15^, thereby quantifying simultaneously the dynamics, size and distance between chromatin blobs in space and time. These results indicated a strong relationship between the flow magnitude of blobs (i.e. their dynamics) and their local density. Notably, the blob dynamics appear enhanced in regions of high blob density and conversely isolated blobs in chromatin-void regions appear less mobile.

**Figure 1:**
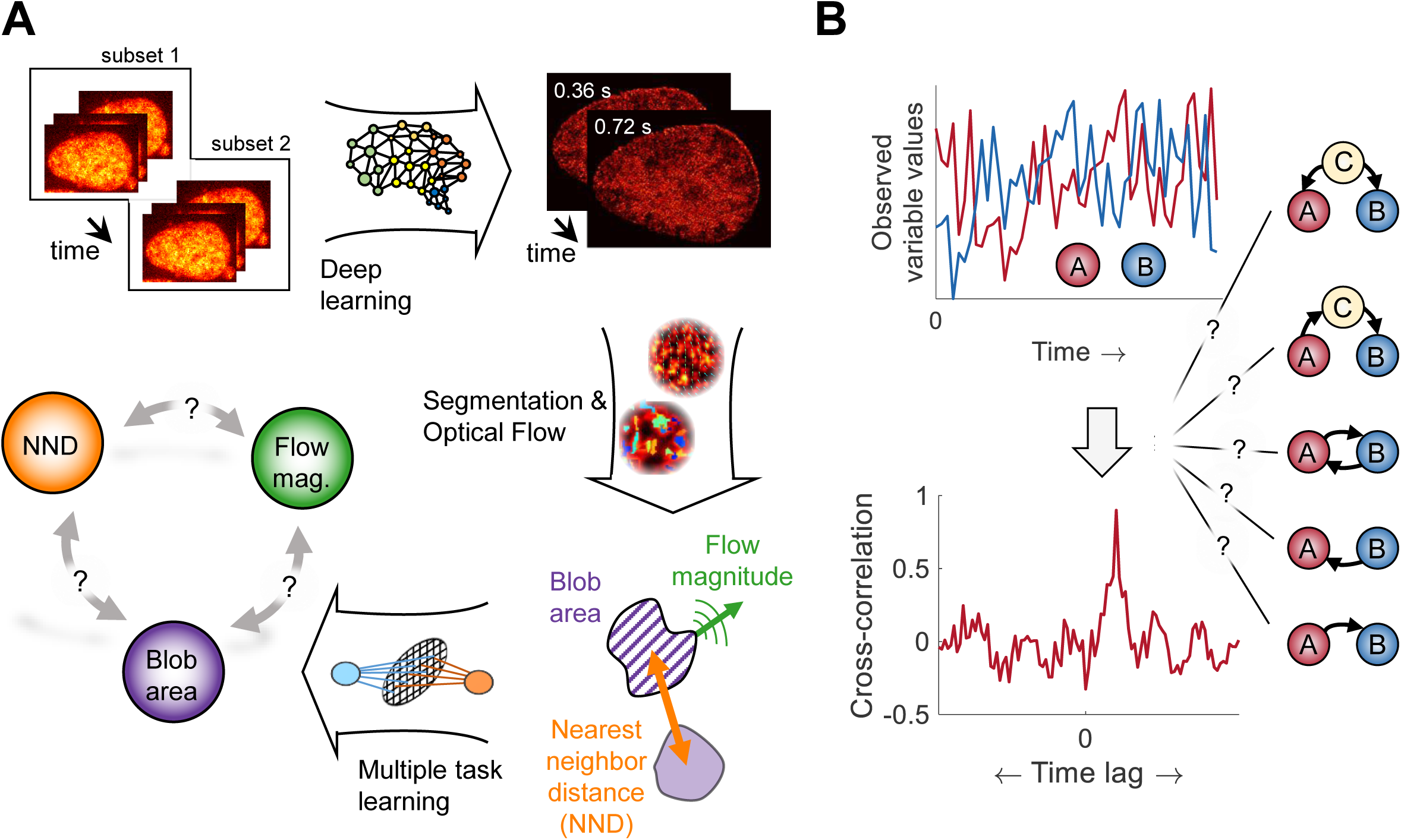
From super-resolution images of fluorescently labeled chromatin to Granger-causal inference between its structure and dynamics. **A)** Human osteosarcoma U2OS expressing H2B-PATagRFP cells were imaged. Deep-PALM combines the predictions of a deep learning algorithm from a subset of 12 images to reconstruct a super-resolved image of chromatin with a temporal resolution of 360 ms at 63 nm spatial resolution ^21^. Segmentation of chromatin blobs and Optical Flow analysis allows ascribing a nearest neighbor distance (NND), blob area and flow magnitude to each blob in each frame individually. Finally, to infer a Granger-causal relationship between these characterizing parameters, multi-task learning is employed. **B)** Whether two observed stochastic variables A and B are operiationally related may be tested by computing a cross-correlation between those variables. The time lag between the variables can be inferred from the absolute maximum value of the cross-correlation curve. However, such a correlation analysis cannot reveal whether the relationship involves causation and in which direction causality is present. Some of the simplest scenarios in a biological context are depicted (from top to bottom): A causes B; B causes A; A and B are in a feedback loop; A causes B indirectly via C; C is the common cause of both A and B.

This and other studies resulted in complex high-dimensional datasets, which reflect stochastic and heterogeneous quantities characterizing chromatin in space and time. In order to allow inferences of relationships in such data, correlations between variables are commonly investigated (Figure 1B). While this is a valid approach, one has to be aware of its limitations ^23^. In particular, a correlation between variables does not directly imply a causal relationship and in the case that a direct causal relationship indeed exists, it is not clear in which direction. Furthermore, the causality might be indirect, transferred via one or multiple unobserved constituents of a biological pathway (Figure 1B). The inference of a causal relationship is thus more powerful than the observation of a correlation, especially in complex systems such as the genome. A methodology to infer causal paths between the chromatin structure, dynamics and ultimately function is thus highly desired, but difficult to obtain. The exploration of true causal relationships would indeed require the complete knowledge of the system, in the sense that every possible influence on genome structure and dynamics (proteins, nuclear morphology, etc.) should be mapped at sufficiently high resolution in three-dimensional space and time. Although approached using simple organisms, absolute complete knowledge of a living system is still virtually impossible^24^ as one can hope to simultaneously capture only a few out of the vast spectrum of parameters at best.

The concept of Granger-causality ^25^ circumvents this issue by inferring Granger-causal relationships only among a subset of experimentally observable variables. In particular, the analysis of Granger-causality is based on the identification of essential variables (the cause) to predict a target variable (the effect). While both correlation as well as causality analyses suffer from the presence of unobserved variables, identification of causal relationships even among the subset of observable variables in the system can, however, indicate which variable is cause and which is effect (via a possibly yet to be discovered pathway).Here, we adapt the abstract concept of Granger-causality to the analysis of Granger-causal relationships between chromatin dynamics and structure at the nanoscale using the unique data set of our recent chromatin live-cell super-resolution imaging based on deep learning (Deep-PALM) ^21^. We use the chromatin blob flow magnitude as a measure of local chromatin dynamics and the blob nearest neighbor distance (NND) as well as the blob area as structural parameters. Our analysis revealed a unidirectional Granger-causality from flow magnitude to both blob NND and blob area, in pixels representing about 10 to 20% of the nuclear volume. We discuss these findings in the light of the theory of active polymers and we further reason that the pervasive activity of Active Effectors on the chromatin fiber may qualitatively and quantitatively explain (i) why chromatin blobs exist and (ii) how the dynamics may influence their structure with respect to inter-blob contacts.

## Materials and Methods

### Cell Culture

U2OS expressing H2B-PATagRFP cells were cultured in DMEM (with 4.5 g/l glucose) augmented with 10% fetal bovine serum (FBS), 100 μg/ml penicillin, 2 mM glutamine, and 100 U/ml streptomycin were incubated at 37°C and in 5% CO_2_. Cells were plated 24 hours before imaging on 35 mm Petri dishes with a #1.5 coverslip like bottom (ibidi, Biovalley) with a density of 2×10^5^ cells/dish. Shortly before imaging, the growth medium was replaced by Leibovitz’s L-15 medium (Life Technologies) supplemented with 10% FBS, 100 μg/ml penicillin, 2 mM glutamine and 100 U/ml streptomycin.

### PALM Imaging

The Deep-PALM imaging conditions are described in our recent publication ^21^. Briefly, a fully automated Nikon TI-E/B PALM (Nikon Instruments) microscope equipped with incubator was used for live cell imaging. NIS-Elements software was used for acquiring the images at 30 ms per frame. PATagRFP was illuminated using a laser line of 561 nm (∼50-60 W/cm^2^ at the sample along with the 405 nm laser line for photo-activation (∼2-2.5 W/cm^2^ at the sample). Excitation wavelengths were merged into a TIRF oil immersion objective (1.49 NA, 100x; Nikon). The same objective was used for collecting the fluorescence emission signal and spectrally filtered by a Quad-Band beam splitter (ZT405/488/561/647rpc-UF2, Chroma Technology) with Quad-Band emission filter (ZET405/488/561/647m-TRF, Chroma). Then the signal was recorded on an EMCCD camera (Andor iXon X3 DU-897, Andor Technologies) with a pixel size of 108 nm.

### Deep-PALM analysis and image processing

Super-resolution images were obtained using a custom-trained convolutional neural network (CNN, ref. ^26^) with an effective pixel size of 13.5 nm. Individual chromatin blobs were segmented using an adapted marker-assisted watershed algorithm and the blob centroid position and area were computed. Using additionally Optical Flow to reconstruct flow fields of chromatin ^14,15^, each blob was ascribed three parameters: its area, its nearest neighbor distance (NND) and its (instantaneous) flow magnitude. In order to retrieve a gridded representation of all variables, the variables were subsequently interpolated onto a 5-fold down sampled pixel grid, resulting in an effective pixel size of 67.5 nm. Details on the super-resolution reconstruction as well as on the segmentation and dynamic analyses can be found in ^21^.

### A framework for the inference of Granger-causality in spatio-temporal data

Granger-causality is assessed between a target variable *Y* and an input variable *X*_1_, potentially conditioned on one or several common variables *X*_2_, *X*_3_, etc. of the system. We observe each variable at each grid point *l* (i.e. pixel) across the entire nucleus and at each time point *t*. The target variable at location *l* is denoted 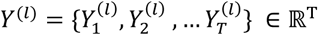, where *T* is the number of time points. Here, *T* = 166, covering a total of ∼ 60 *s* with a time resolution of Δ*t* = 360 *ms*. Similarly, 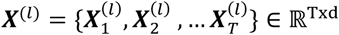, where 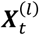 is a tupel of the cause variable 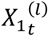 and the *d* − 1 conditional variables 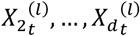, etc. at location *l* and time *t. d* thus denotes the number of input variables, including the potential cause. As stated in the main text (Figure 2A), testing for Granger-causality involves modeling of the target *Y* using the lagged variables ***X*** (ref. ^25^). The base model excludes the variable *X*_1_ for which Granger-causality is tested:

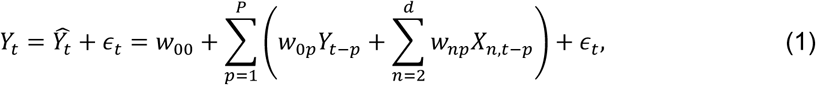

where *Ŷ*_*t*_ is the model prediction of the true target *Y*_*t*_, *ϵ*_*t*_ is a residual noise term, the matrix *w*_*ij*_ denotes the coefficients of the base model and *P* is the maximum time lag considered. The full model contains all explicit and conditional cause variables *X*_1_, …, *X*_*d*_:

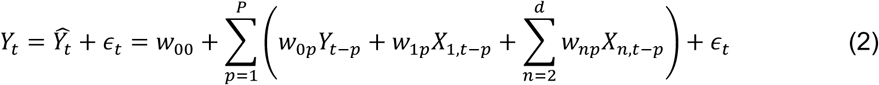

**Figure 2:**
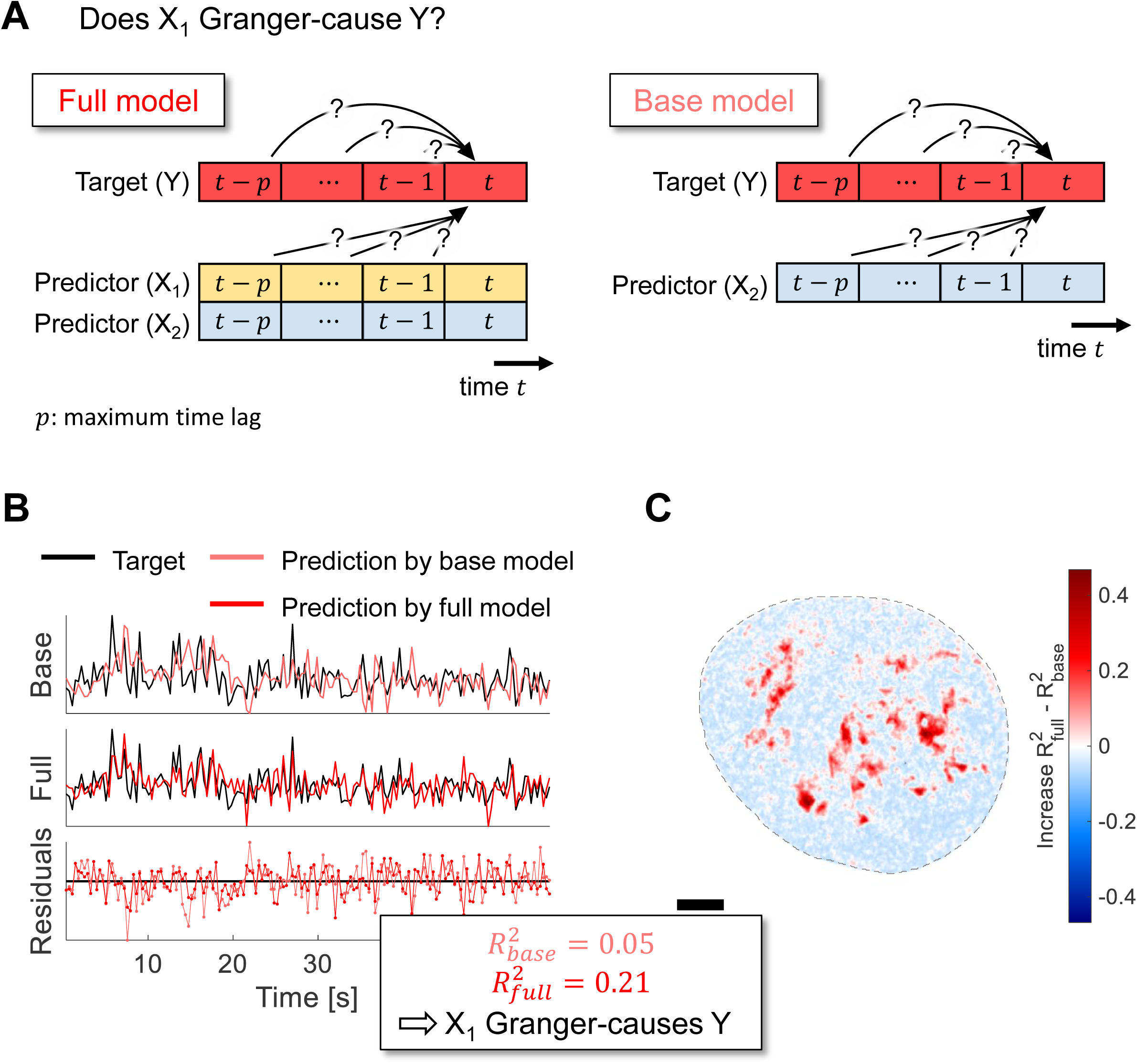
Inference of Granger-causality. **A)** The value of a target variable *Y* at time *t* may be determined by its own past and the past values of other variables in the system *X*_1_ and *X*_2_. The full model (left panel) consists of a linear relationship between the values of *Y, X*_1_ and *X*_2_ from the time point *t* − *p*, where *p* indicates the maximum time lag considered, to time point *t* − 1. In contrast, the base model takes only variables *Y* and *X*_2_ into account (right panel). **B)** Both models are independently optimized to model the target *Y* and their prediction accuracy is assessed by computing the adjusted *R*^2^ via their residuals. If and only if the adjusted *R*^2^ value of the full model is positive, higher than of the base model and the Diebold-Mariano statistic is significant (Materials and Methods), *X*_1_ is said to Granger-cause *Y*. In the depicted example, *Y* is the blob NND at a chosen pixel within the nuclear interior. All time traces have been scaled to mean zero and unit variance. **C)** An exemplary map across a nucleus showing the difference between the adjusted *R*^2^ values of the full and the base model. The target variable is the blob NND. Positive values (red) indicate that the variable under consideration (here *X*_1_) considerably improves the modeling of *Y* and therefore Granger-causes *Y*, while negative values (blue) indicate that no Granger-causality can be detected during the time of the observation.

Note that the second sum runs from 2 to *d* in the base and the full model and the contribution of the cause-variable *X*_1_ is written explicitly in the latter. Granger-causality from *X*_1_ to *Y* is present if the predictions *Ŷ* of the full model (including the past value of *X*_1_) are significantly better than those of the base model. Quantitatively, we assess the significance in two complementary ways. First, we use the Diebold-Mariano test ^27^ to evaluate if the predictions of the two models are significantly different. Then, we compute the adjusted *R*^2^ value to account for the varying number of coefficients in the base and full model ^28^:

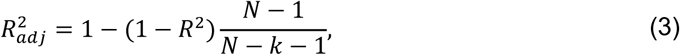

where *N* is the total number of samples, *k* is the number of parameters in the model and *R*^2^ is the coefficient of determination:

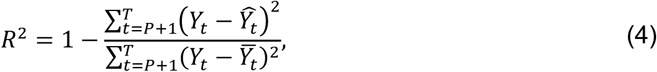

with 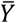 denoting the time average. Pixels at which the adjusted *R*^2^ of the full model is negative or the Diebold-Mariano test does not indicate a significant difference between the prediction errors of full and base models are marked as non-causal.

### Multi-task learning

We aim to solve regression problems of the form of equations (1) or (2) in order to find the matrix *w*. This can be done for each pixel individually by minimizing a loss function *Ψ*:

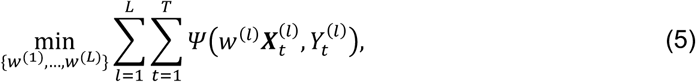

where *L* is the number of locations. However, it was shown previously that chromatin dynamics ^14,15^ as well as the chromatin structure ^21^ are spatially correlated. It is therefore reasonable to exploit the fact that different locations share a similar behavior. We therefore use multi-task learning, following an earlier approach to fit linear regression models to spatially correlated time-series in the geoscience domain ^29^. In this work, the authors use an alternative structure optimization (ASO, ref. ^30^) method in order to simultaneously learn a shared low-dimensional representation among the tasks. In particular, the weight matrix *w* is split into two parts: *w* = *u* + *ν*Θ, where *u* is a weight matrix in the original *d*-dimensional space, *ν* is a weight matrix in the shared low-dimensional space and Θ is a parameter matrix with orthonormal row vectors (ΘΘ^′^ = **1**). The ASO multi-task learning optimization can be thus expressed as

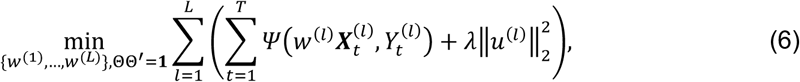

With the regularization term 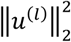 and a regularization parameter *λ*. This term penalizes weight differences between the high-dimensional and low-dimensional space, parametrized by Θ. A memory-limited Broyden– Fletcher–Goldfarb–Shanno (L-BFGS) algorithm ^31^ is used to optimize equation (6). Here, we use a maximum time lag of 3.6 s (10 data points) for the regression, which allows for the incorporation of reasonably long-time scale in the inference, while the number of optimizable parameters remains feasible (given time series consisting of 166 data points). The number of parameters equals the number of variables in the system times the number of time points (3 variables times 10-time lags in this case). Further details on the regression can be found in ^29^.

The multi-target learning algorithm uses the information of spatially related processes and therefore enhances its prediction accuracy ^29^. The resulting weight matrix *w* is used to construct the model predictions 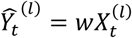, which are used to evaluate the prediction performance of the base and full model in terms of 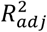.

## Results

### A framework to infer Granger-causality between chromatin dynamics and structure

The probability that a true causal relationship with all of the engaging components can be established in a biological experiment is very small – usually, only a subset of all potential direct, indirect or hidden variables can be observed (see the indirect causal relationships via an unobserved component ‘C’ in Figure 1B). We decided therefore to employ the concept of Granger-causality to deduce causal relationship between chromatin structure and dynamics ^25^. Briefly, a time series *X*_1_ is said to Granger-cause *Y*, given another observable time series *X*_2_ inherent in the system, if the past values of *X*_1_, *X*_2_ and *Y* can predict *Y* at the current time *t* better than *X*_2_ and *Y* alone. This definition can, of course, be extended to include additional observable time series in the system apart from *X*_2_ if such data becomes available. However, since our data set comprises three variables of interest, we stick to one target, one predictor and one conditional variable for simplicity. The concept is illustrated in Figure 2A. A basic principle of causality is that there is no instantaneous influence of the cause to the effect. Instead, the value of *Y* at time *t* can be modeled as a linear superposition of past values of *X*_1_, *X*_2_ and *Y* up to a maximum time lag considered (for details on the model regression strategy, see Materials and Methods). This constitutes the ‘full’ model, in which all observed system variables are taken into account (Figure 2A; left). In contrast, the ‘base model’ uses only past values of the common variables *X*_2_ and *Y* (Figure 2A; right). The prediction accuracy of both models is then evaluated using the residuals between the predicted and true values of *Y* at time *t* (Figure 2B) and quantified using the adjusted *R*^2^ value (Materials and Methods). Additionally, a Diebold-Mariano test is used to consider only significant differences in the prediction accuracy of the two models (Materials and Methods). The full and base model are evaluated at all pixels within the nucleus. Pixels at which the adjusted *R*^2^ value of the full model is higher than of the base model indicate pixels at which a Granger-causal relationship can be detected (Figure 2C).

While this framework is not able to address head-on the true causality for a system as complex as a human cell (the current state-of-the-art experimental data do not allow to do so), it is able to infer relative causalities (Granger-causalities) among the set of accessible parameters. The detection of Granger-causality thus indicates that some path is present from one variable to another such that the latter appears to arise as a consequence of the former. Note that inference of Granger-causality adds information, compared to a correlation analysis, of the direction of influence and hence allows to classify two variables as cause and effect. However, it is not possible to directly analyze which molecular constituents of the system are involved and which pathway is responsible for the observed causality. The framework presented here therefore represents a first step towards a more extensive inference of causal relationships in the highly complex context of chromatin in space and time. Moreover, it opens avenues to truly identify and decipher the mechanisms responsible for dynamic and functional chromatin organization in the future.

### Chromatin dynamics act upstream of blob density

We imaged H2B-PATagRFP in live human bone osteosarcoma (U2OS) cells for up to 60 seconds using Deep-PALM, a live chromatin super-resolution technique^21^. H2B is one of the 4 core histones found in every nucleosome, and imaging H2B-PATagRFP thus constitutes a way to follow the motion of chromatin in a nucleus.

We analyzed a series of 166 super-resolved frames of dynamic chromatin with a time resolution of 360 ms, governing ∼ 60 seconds, and a spatial resolution of 63 nm. Segmentation of the spatially heterogeneous H2B signal identified ∼ 10,000 chromatin ‘blobs’ at any time in a U2OS nucleus (Figure 1A). The blobs appear to result from the dynamic and stochastic association of a number (< 30) of nucleosomes in groups^21,22^. Comparison of super-resolution images to polymer simulations suggested that blobs can be identified as sub-TADs in the time-average limit, while blobs assemble/dissociate on the time scale of about one second^21^. However, it is to date unknown which (functional) mechanism determines their formation and characteristics. Chromatin blobs were experimentally characterized in terms of their area, axial dimensions (45 to 90 nm wide elongated shape) and their nearest neighbour distance (NND) between each other^21^. Analysis of the apparent bulk chromatin motion across the image series using Optical Flow ^14,15^ allowed moreover to ascribe an instantaneous flow magnitude (velocity) to each blob (Figure 1A).

We applied the framework introduced above to these three parameters characterizing chromatin dynamics (the instantaneous flow magnitude) and organization (the NND between blobs and the blob area). The three parameters could potentially exhibit causal relationships in any direction and also participate in feedback loops (Figure 1B). We therefore tested all possible combinations for Granger-causal relationships. Pixels within an exemplary nucleus at which Granger-causality was detected are marked by the respective color of the cause (Figure 3A). In general, when Granger-causality was detected, it was observed in no more than 20% of the nucleus suggesting a picture where stochasticity dominates order. Remarkably, the flow magnitude appears to act upstream and determine the NND and blob area, while the inverse relationship was hardly ever observed (<1%). This indicates that chromatin dynamics is a key determinant of chromatin organization at the nanoscale. Such Granger-causal relationships could be demonstrated in essentially non-overlapping areas of the nucleus for each parameter. That dynamics are Granger-causal for the blob density was observed in large micrometer-spanning, connected regions and mostly in the nuclear interior (Figure 3B). In contrast, dynamics was found to be a Granger-cause for the observed blob area in smaller regions and rather closer to the nuclear periphery (Figure 3B). The NND and blob area further appear connected in a feedback loop in pixels scattered throughout the nucleus, suggesting they are Granger-cause and effect of each other. A causal loop diagram shows the direction of the observed Granger-causalities and highlights that these Granger-causal relationships could be demonstrated in ca. 10% of pixels in every case. However, it is possible that the diagram applies to chromatin in general but can be demonstrated only when and where order supersedes stochasticity for a given parameter (Figure 3C).

**Figure 3:**
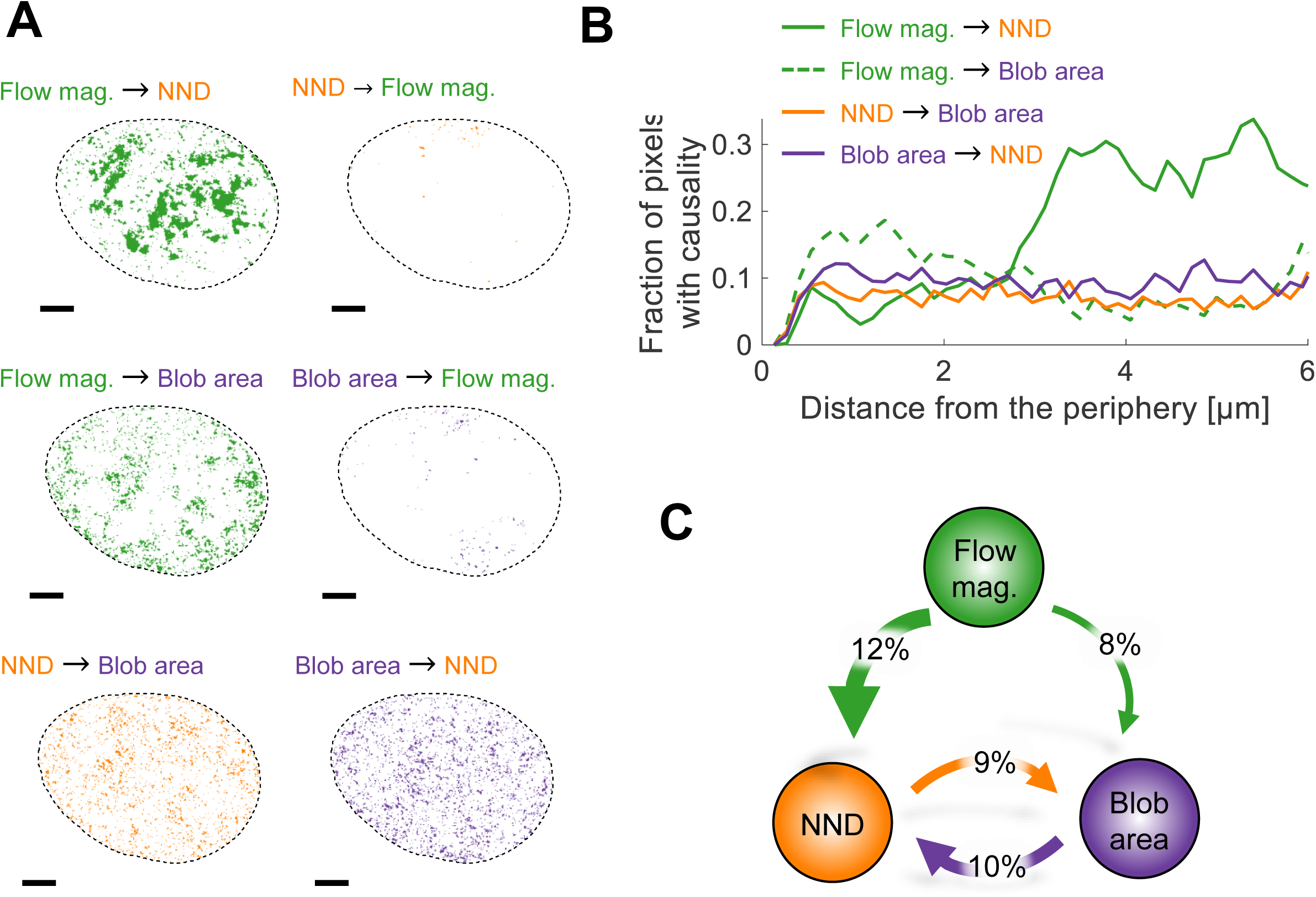
Chromatin flow magnitude Granger-causes chromatin structure. **A)** All combinations between potential Granger-causal relationships of the system variables were tested. An exemplary nucleus is shown for which pixels are colored according to the cause. The flow magnitude (green) is shown to mainly cause structural parameters to vary, while the inverse was barely observed. Scale bar is 3 µm. **B)** The fraction of pixels for which Granger-causality was observed is shown in dependence of the distance to the nuclear periphery. **C)** A loop diagram summarizing the Granger-causal relationship between the flow magnitude, NND and blob area. Percentages correspond to the average frequency across the data set at which pixels were detected with a Granger-causal relationship in the indicated direction, relative to the nucleus size. Percentages < 2% were omitted for clarity.

For the cases, in which causal relationships could be identified, we analyzed the (temporal) regression weights for the target, predictor and conditional variable nucleus-wide (**Error! Reference source not found**.A). The temporal weights indicate the relevance of a variable at time *t* − Δ*t* to predict the target variable at time *t*, where Δ*t* is a time lag. Concomitantly, the temporal cross-correlation between target and predictor or target and conditional variable, respectively, is shown for the first 10-time lags (3.6 seconds; **Error! Reference source not found**.B). In general, blob NND and area exhibit an inverse relationship in both the regression weights and the temporal cross-correlation (see below). The influence of the flow magnitude on the regression is largest when the flow magnitude is identified as the cause of a structural parameter and almost negligible otherwise, as expected (**Error! Reference source not found**.A). In contrast, the cross-correlation between flow magnitude and either NND or area (**Error! Reference source not found**.B) differs from zero for the considered time lags irrespective of the identified causality. This demonstrates that a causal relationship between two variables does not necessarily exist, even when the parameters are correlated. It is likely that the correlation is established via one or multiple unobserved factor(s).

We next examined whether individual parameters showed any noticeable deviations in regions in which Granger-causality could be demonstrated as compared with the rest of the nucleus. The dynamics was found to be higher and blobs were, on average, closer in regions in which the flow magnitude was shown to act upstream of the blob density (Figure 4A-C). This finding is in line with a correlation approach which previously found that blobs with close neighbors are on average more dynamic than blobs further away from each other ^21^. Here we extend this observation by noting that blobs arise with a higher density actually *as a consequence of* locally elevated dynamics. In contrast, the flow magnitude appears overall similar for pixels in which the flow magnitude could be demonstrated to be a cause for the blob surface area, as compared with the rest of the nucleus (Figure 4E). In these restricted regions however, blobs tended to be smaller and blob density was clearly lower on average (Figure 4F, G), an inverse trend as compared with regions in which flow magnitude was shown to act upstream of blob density. Regarding the Granger-causal loop involving only the NND and blob area, no significant bias in any parameter under consideration could be evidenced (**Error! Reference source not found**.). Whenever a Granger-causal relationship from chromatin dynamics to a structural parameter was found, the temporal cross-correlation was slightly but significantly enhanced at long time lags (Figure 4D, H), indicating that the structure-dynamics coherence was sustained over an extended time in these regions. It should be noted that the size of chromatin blobs in relatively chromatin-void regions is likely to be well captured in our analysis due to a good signal-to-noise ratio, whereas in denser chromatin regions multiple small but close chromatin blobs might be detected as merged into a single bigger blob. This artificially increases the measured blob surface area, while the NND is more robust to a blob merging in regions with high blob density (**Error! Reference source not found**.). The following discussion and modeling are therefore based only on the two more reliable parameters, blob dynamics and NND as a proxy for blob densities.

**Figure 4:**
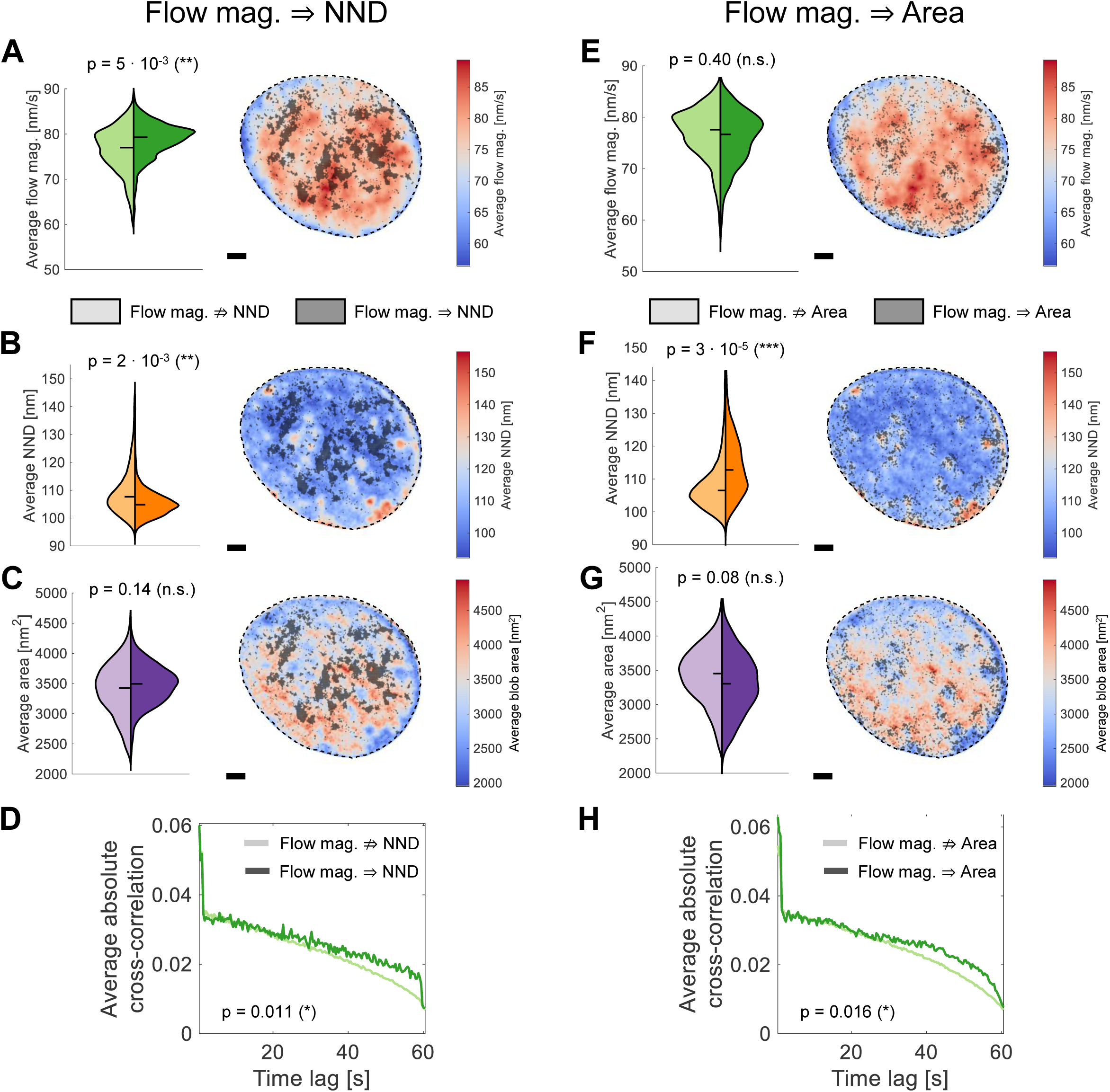
Enhanced dynamics and dense chromatin in regions with Granger causality. **A)** The time-averaged flow magnitude across nuclear regions in which no Granger-causality was detected (bright color in the violin plot and the map on the right) and in regions in which the flow Granger-causes the NND of blobs. Scale bar is 3 µm. **B-C)** The average NND and blob area in the same manner as for A). **D)** The average absolute cross-correlation between flow magnitude and NND for pixels in the two regimes (with/without Granger-causality). The correlation is significantly enhanced in the long-time limit for regions in which Granger-causality is detected. **E-H)** As for A-D) in the case of a detected Granger-causality from the flow magnitude to the blob area. Statistical significance was assessed using a Wilcoxon rank sum test. The shown p-value is the median from 250 tests on sub-sampled data to avoid reporting a significance due to the large sample size alone.

In summary, we show that blob dynamics and blob structural parameters are strongly connected and chromatin organization appears to arise as a consequence of chromatin dynamics. Such a cause-and-effect relationship can only be demonstrated in restricted areas of the nucleus, probably because stochasticity overrides deterministic mechanisms everywhere else. Strikingly, higher dynamics was found to be a cause for increased blob density (Figure 4; left column). Since biological experiments can only give access to a small subset of variables and influences, it should be kept in mind that the described causal relationships refer to the concept of Granger-causality. It is probable that the chromatin dynamics are not the true cause, but that dynamics are in fact mediated by other mechanisms such as ATP-driven processes, entropic considerations, etc. Furthermore, the dynamics-structure Granger-causality described here can be coupled via one or several unobserved factors. Below, we discuss a possible polymer-theoretical approach and different biological processes that (i) can induce chromatin dynamics beyond passive thermal or entropic contributions (ii) could potentially give rise to the formation of transient chromatin blobs and (iii) could potentially explain how chromatin dynamics can shape the transient chromatin structure on the length scales of blobs.

## Discussion

The inference of (Granger-) causality in complex systems provides a powerful tool to infer mechanisms from observations. Below we show that the results of our Granger-causality analysis are consistent with the theory of active semiflexible polymers. We discuss the major findings of simulations and analytical descriptions in a qualitative, intuitive way and describe implications of this theory for the organization and dynamics of chromatin. Finally, we assess if and under which circumstances chromatin blobs can be caused by an activity-induced motion of chromatin.

### The theory of active semiflexible polymers could explain the existence of chromatin blobs

At the root of active polymers are active particles (self-propelling particles, dipolar motors, enhanced diffusion, etc.). These active particles can be either incorporated as part of the polymer itself (several monomers of the chain are active) or as part of the fluid surrounding a passive polymer (Figure 5A). These active particles are typically Active Effectors (see introduction) which provide a local influx of energy by hydrolysis of ATP and associated chromatin remodeling or simply by inducing deformation of the chromatin fiber upon binding. Variations between models yield qualitatively rather similar results ^32^ and since Active Effectors can act on the chromatin fiber in various ways, we shall neglect model-specific differences in this discussion.

**Figure 5:**
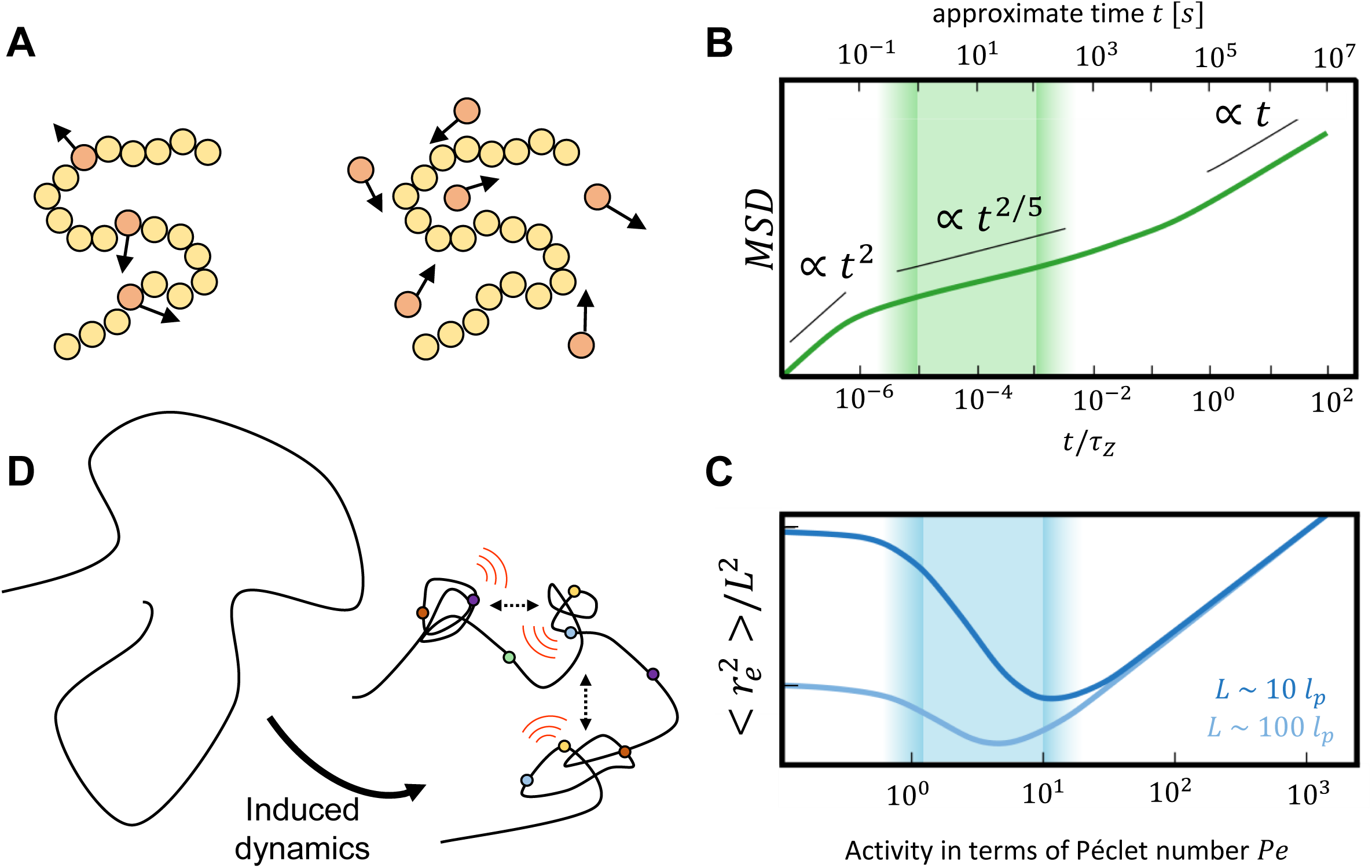
Dynamical and structural properties of active semiflexible polymers. **A)** Active polymers can be described as polymers consisting of a mixture of passive (yellow) and active (red) monomers (left). Alternatively, active polymers are modeled as passive polymers embedded in a bath of active particles (right). Mathematically, a colored noise term can be included in the equations of motion to model activity ^32^. **B)** The mean squared displacement (MSD) of a linear active polymer with ***L* ∼ 10**^**5**^ ***l***_***p***_ at ***Pe* = 20**, subject to hydrodynamic interactions, is illustratively shown versus time lags in terms of the Zimm time ***τ***_***Z***_ of a passive polymer (adapted from ^50^). The upper x-axis shows an approximate mapping to absolute time in seconds (Supplementary Note 1) and the shaded area denotes the experimentally accessible time scale. The straight black lines serve as a guide to the eye to identify the different scaling regimes. **C)** The mean squared end-to-end distance 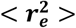 is shown illustratively for two polymers of length ***L* ∼ 10**^**1**^ ***l***_***p***_ (dark blue) and ***L* ∼ 10**^**2**^ ***l***_***p***_ (light blue) over the Péclet number ***Pe*** (adapted from ^33^). The shaded area denotes the biologically relevant regime of the Péclet number (Supplementary Note 1). Ticks along the y-axis indicate one order of magnitude. **D)** According to the theory and simulations of active semiflexible polymers, dynamics induced by a host of chromatin players in distinct classes (i.e. polymerases, chromatin remodelers, topoisomerases, and HMG proteins) (colored dots) can stochastically collapse of loops into chromatin blobs and enhance blob-blob interactions. This results in locally increased chromatin blob density and reduced blob nearest neighbor distances. The growth/shrinkage of blobs which result from the preceding activity along the chromatin fiber could possibly be tracked for isolated blobs in three dimensions.

Rather expectedly, simulations and analytical approaches of active semiflexible polymers reveal that the diffusion of active polymers is enhanced compared to their passive counterpart ^33–37^. The scaling behavior of the mean square displacement transits from a ballistic motion at very short time scales due to the particle propulsion to a subdiffusive regime at intermediate times and a diffusive regime at long time scales ^38^ (Figure 5B). Furthermore, the relaxation time of polymers, i.e. the time needed to recover to the initial conditions after applied stress, decreases with increasing activity ^33^. In line with this observation, an active polymer also exhibits enhanced conformational fluctuations ^37^ and appears more flexible. In particular, the effective persistence length (the length over which a polymer is approximately stiff) of active polymers is decreased compared to their passive counterparts as a consequence of hairpin-formation ^34^.

The dependence of the relaxation time on the activity gives rise to interesting conformational properties of active polymers. The dimensionless Péclet number *Pe*, which is defined as the ratio of the (activity-induced) advection and (passive) diffusion rate in a system (Supplementary Note 1), is commonly used to denote the activity strength in active polymers. For moderate activities as reflected by a low value of Péclet number *Pe*, the polymer experiences a significant shrinkage in terms of its means square end-to-end distance ^33^ and its radius of gyration ^34^. For very large activities, it swells monotonically (Figure 5C). Shrinkage was also described in terms of a coil-to-globule-like transition upon increasing activity ^39^, which might hint towards a reason why chromatin is well described by a fractal globule^40,41^ with many-body contacts ^42^ that moreover displays anomalous diffusion ^43^.

The activity component in these models thus causes (i) enhanced conformational flexibility and diffusion with an intermediate subdiffusive regime and (ii) a shrinkage of polymers in specific regimes, which are determined by the activity and polymer characteristics such as its total length and its persistence length. We propose that the previously observed chromatin nanodomains / blobs ^21,22,44^ might be formed due to the local activity of various Active Effectors and that at least aspects of the dynamic and structural behavior of chromatin blobs can be captured by the theory of active semiflexible polymers (Figure 5D). This notion fits well with the findings that the blob dynamics and NND influence each other over space and time and that chromatin dynamics is at the root of blob organization (Figure 3; Figure 4). Below, we consider biologically relevant and experimentally accessible time and length scales to evaluate if the theory of active semiflexible polymers can be united with experimental findings.

### Experimentally and biologically relevant time and length scales of chromatin blob formation

Chromatin blobs were mainly observed on the length scale of tens ^21,22,44^ to hundreds ^45^ of nanometers. While most studies employed super-resolution microscopy on fixed samples, only two studies gave access to high-resolution chromatin dynamics using Structured Illumination Microscopy ^45^ and Deep-PALM imaging^21^. The time resolution in the latter was 360 ms, while images were acquired for up to 60 s. The experimentally accessible time range is thus in the order of 10^0^ − 10^2^ seconds. The simulated time is estimated in terms of the Zimm time τ_*Z*_, which is in the order of 10^5^ *s* (Supplementary Note 1). Mapping this time approximately to the simulated time scale of the dynamics of active polymers shows that only the subdiffusive regime is experimentally observable (shaded area in Figure 5B), which is in line with the subdiffusion of chromatin also measured in experiments ^11,14,16,21,46^.

Considering the Péclet number *Pe* (defined as the ratio of advection and diffusion rate; Supplementary Note 1), an upper bound on the diffusion constant of chromatin *in vivo* was estimated in the order of *D* ∼10^−3^ *μm*^2^/*s*^α^ (ref. ^14^). As an example for activity induced motion, the speed of loop extrusion of SMC proteins such as condensin in yeast or cohesin in humans was measured *in vitro* as ∼ 0.5 *kb*/*s* (ref. ^7,8^). The Péclet number can thus be estimated in the order of 1 - 10 for chromatin (Supplementary Note 1; shaded area in Figure 5C), demonstrating that the induced activity can indeed contribute to condensing chromatin locally. However, there might be exceptions beyond the estimated time and length scales at which the Péclet number might be as large as to cause swelling of polymers (Figure 5C).

### The treatment of active circular polymers and polymer-polymer interactions can recapitulate experimental observations

The considerations above demonstrate that activity-induced chromatin blob formation is conceivable. The observation that changes in the nearest-neighbor distance of chromatin blobs can be frequently traced back to be caused by their dynamics (Figure 4A-C) further lends support to the theory. Considering pixels in which the flow magnitude influences the area of isolated blobs (Figure 4E-G), the area fluctuations could reflect a growth/shrinkage of blobs according to the preceding dynamics (Figure 5D). The fact that those blobs appear far from their neighbors opens up the possibility to track these blobs in three dimensions to verify this hypothesis. Nevertheless, a direct translation of the theory of active semiflexible polymers remains difficult due to the multitude of influences on chromatin that exist *in vivo*. A few additional mechanisms are treated in the literature. Considering structural elements of chromatin such as TADs, sub-TADs as well as chromatin loops such as those that can be extruded by SMC complexes ^7,8^, one should take into account that chromatin blobs likely correspond to sub-regions of quasi-circular, not of linear polymers. However, the results for linear active polymers are qualitatively transferable to circular ones ^37^, strengthening the hypothesis that blobs could correspond to loops or sub-TADs ^21^.

Activity was independently shown to enhance the looping probability of chromatin segments ^47^. Looping may be further stabilized by DNA bridging factors and/or SMC proteins at the loop bases ^48^ as well as a host of transcription factors operating as dimers. Of note, there are hints that crowding may further promote blob formation in a crowder size and concentration-dependent manner ^49^, and taking into account hydrodynamic interactions can lead to even further shrinkage of polymers^50^.

Interestingly, a simulation involving ensembles of active semiflexible polymers showed that the different polymers get closer with increasing activity ^35^. This computational result is in line with our observation that regions in which a high blob density is shown to depend on chromatin dynamics are also regions where dynamics is more elevated (Figure 4). Altogether it appears that activity can promote blob-blob interactions by simultaneously enhancing blob mobility and decreasing the distance between blobs to make chromatin locally more compact and disordered (Figure 5D).

## Conclusions

Using a framework to infer Granger-causal relationships between spatio-temporal variables derived from a previous whole-chromatin live super-resolution imaging study ^21^, we analyzed if and how chromatin dynamics and organization influence each other. Within a subset of simultaneously observable variables of a system, this framework allowed us to pinpoint directed Granger-causal relationships among parameters beyond the more conventional description of correlations alone. Within the limitations of our data set, we found that dynamics can be considered as a cause of structural parameters, and in particular that locally elevated chromatin dynamics causes blobs to be closer to each other. This is a rather counter-intuitive result as high chromatin density is commonly associated with closed chromatin, in which reduced chromatin density is expected due to increased constraints on DNA. Our results suggest that closed chromatin is in fact a very active environment as further supported by the fact that a number of Active Effectors are known to be key determinants of closed chromatin assembly and function^10,51^. In addition, active processes have been shown experimentally ^52,53^ to drive coherent motion of chromatin ^15,16^.

To gain further insights into the existence of possible spatio-temporal causal relationships, the presented analysis may be extended to include the influence of variables at neighboring pixels. This is particularly important as chromatin blobs naturally move from frame to frame. However, since the number of regression parameters scales with the number of neighboring pixels (3 parameters x 10 time lags x 4 or 8 neighboring pixels = 120 to 240 parameters), the current length of the time series (166 data points) does not allow for reliable inference of causality. Further enhancing the time resolution of chromatin super-resolution imaging and circumvention of photobleaching for longer acquisition can alleviate this restriction in the future.

We demonstrate more broadly that the theory of active semiflexible polymers has the potential to explain the experimentally observed characteristics of chromatin blobs on biologically relevant scales, and can further provide an intuitive explanation for the observation that increased blob mobility can locally co-exist with dense chromatin. Blobs as we are able to observe them may nevertheless arise as a result of further stabilization by bridging and/or cross-linking factors. To probe this theory more explicitly, the analysis presented here may be carried out in cells which are depleted in certain key factors such as SMC proteins or ATP. Especially in ATP-depleted cells, we expect a global loss of causality.

Our analysis altogether reveals that chromatin dynamics is a key determinant of genome organization in nuclear space. However, such Granger-causality could be demonstrated only in restricted areas of the nucleus that are largely non-overlapping for distinct combinations of parameters. The identification of Granger-causal relationships *per se* throughout the nucleus indicates that (multiple) deterministic molecular mechanisms exist and constantly remodel chromatin. The sparsity of such Granger-causal relationships, however, also demonstrate that chromatin is governed by stochasticity^54^.

## Acknowledgments

We acknowledge support from the Pôle Scientifique de Modélisation Numérique, ENS de Lyon for providing computational resources and further thank the Institut Rhônalpin des Systèmes Complexe IXXI for supporting us.

## Author contributions

R. B. and H. A. S. designed the project; R. B. designed and carried out the data analysis; R. B. and H. A. S. interpreted results; H. A. S. supervised the project; R. B., G. F. and H. A. S. wrote the manuscript.

