## Supplementary material for "Dynamics as a cause for the nanoscale organization of the genome": Supplemntary

### Revealing causality between chromatin dynamics and organization at nanoscale resolution


R. Barth[1], G. Fourel [2, 3] and H. A. Shaban [4, 5, *]

1: Department of Bionanoscience, Delft University of Technology, 2628 CJ Delft, The Netherlands.

2- Laboratory of Biology and Modelling of the Cell, University of Lyon, ENS de Lyon, University of Claude Bernard, CNRS UMR 5239, Inserm U1210, Lyon, France.

3: Centre Blaise Pascal, ENS de Lyon, Lyon, France.

4: Spectroscopy Department, Physics Division, National Research Centre, Dokki, 12622 Cairo, Egypt.

5: Center for Advanced Imaging, Northwest Building, Harvard University, Cambridge, MA, 02138, USA.


#### Supplementary Note 1: Estimation of time and length scales to match theory and experiment

**Time scale.** The time in **Error! Reference source not found.**B is scaled by the Zimm time $\tau_Z \sim \eta_s R_F^3/(\sqrt{3\pi} k_b T)$, where $\eta_s$ is the solvent viscosity, $k_b T$ is the thermal energy and $R_F \sim \sqrt{N} b$ is the Flory radius of a Gaussian chain with $N$ the number of monomers and $b$ the bond length [1]. We take twice the persistence length $l_p$ of DNA as an estimate for the bond length, where the persistence length is usually estimated between 50 and 200 nm [2], roughly in the order of $l_p \sim 10^2\ nm$. For this discussion, we do not take the whole genome into account, but sub-compartments of chromatin *in vivo,* which could be topologically associated domains (TADs) or loops formed by either bridging or loop extruding factors [3–6]. Loops can be hundreds of kilobases ($\sim 10^2\ kb$) long [7].

Chromatin consists of DNA wrapped around nucleosomes, which form the basic structural element; about 200 bp are wrapped around one nucleosome, taking into account linker DNA of up to 80 bp. In spatial dimensions, the nucleosome is about 10 nm wide and the linear linker DNA is about 30 nm long. Therefore, we consider one stretched out nucleosome to be roughly a few tens of nanometers long. Considering the loop dimensions of $10^2\ kb$, there are in the order of $10^3$ nucleosomes involved, which would measure $\sim 10^4\ nm$ when stretched out linearly, corresponding to about $\sim 10^2\ l_p$. We can thus estimate the Flory radius as $R_F \sim 10\ l_p$.

With the thermal energy of about $k_bT \sim 4\ pN \cdot nm$ and estimates of the viscosity inside the cell nucleus in the order of $\eta_s \sim 10^3\ Pa \cdot s$ [8], we arrive at an estimate of the Zimm time in the order $\tau_Z \sim 10^5\ s$. We can thus approximately convert the simulated time in units of the Zimm time in **Error! Reference source not found.**B to absolute time in seconds and compare with the experimentally accessible time scales.

**Length scale.** Following the considerations above, we recapitulate the order of magnitude for the persistence length $l_p \sim 10^2\ nm$ and the total linear length of chromatin loops *in vivo* $L \sim 10^4\ nm$. The ratio $L/(2l_p)$, which is commonly used in the theoretical treatment of active polymers [9–11], can thus be estimated to be in the range of $L/(2l_p) \sim 10^2$. The two lines shown in **Error! Reference source not found.**C correspond to the cases $L/(2l_p) \sim 10^1$ (dark blue) and $L/(2l_p) \sim 10^2$ (light blue).

**Magnitude of dynamics.** Active polymers are characterized by the dimensionless Péclet number $Pe = UL/D$, which determines, loosely speaking, the ratio between the advection and diffusion rate in a system [12]. Here, $U$ is the linear flow velocity, $L$ is the length scale over which advection occurs and $D$ is the (passive) diffusion constant. For $Pe < 1$, the system is diffusion-dominated, while the system is advection-dominated for $Pe > 1$. To give an order of magnitude of the Péclet number of chromatin flow, one should be able to disentangle purely diffusive and activity-induced flows in living systems. To give a lower bound on the expected Péclet number, we consider the diffusion of chromatin under the influence of ATP-driven processes. Chromatin diffusion coefficients are found to be in the order of $D \sim 10^{-3}\ \mu m^2/s^\alpha$ [13], while purely passive diffusion due to thermal fluctuations is expected to be lower [14]. We next consider the advection rate induced by active processes. Consider for instance transcription elongation by RNA polymerase II, which proceeds at around $U \sim 2\ kb/min$ for $L \sim 100\ kb$ [15] or equivalently $U \sim 5\ nm/s$ for $L \sim 10\ \mu m$. This leads to a Péclet number in the order of $Pe \sim 1$. Another example is the loop extrusion activity by SMC complexes, which proceeds with a speed of $U \sim 0.5\ kb/s \sim 10^2\ nm/s$ [4,6] for about $L \sim 10^2\ l_p \sim 10\ \mu m$, following the considerations above, yielding $Pe \sim 10$. These examples demonstrate a lower bound for the Péclet number in the order $Pe \sim 10^0 - 10^1$ of chromatin flow *in vivo*. This range is indicated as biologically relevant regime as shaded area in **Error! Reference source not found.**C.

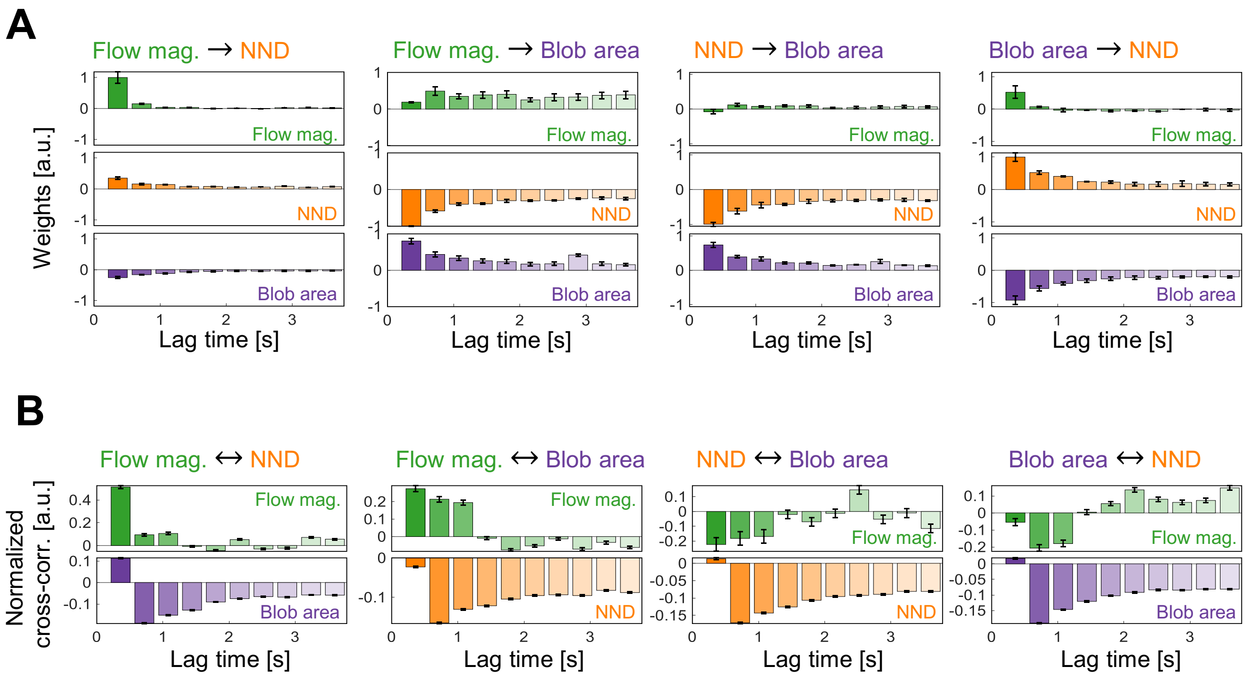


**Supplementary Figure 1: Temporal regression weights and cross-correlation. A)** For each unidirectional Granger-causal relationship, the average (nucleus-wide for those pixels for which a Granger-causal relationship was identified; error bars denote the standard error) regression weight is shown over the first 10-time lags considered (see Materials and Methods). Each predictor, conditional and target variable at time $\boldsymbol{t - \Delta t}$ can contribute to the value of the target variable at time $\boldsymbol{t}$, thus giving rise to 30 (3 variables times 10-time lags) regression weights per pixel. **B)** The normalized average cross-correlation for the 10-time lags between the predictor and target as well as the conditional and target variable, respectively (correlation between target and target variable was omitted since the analysis focuses on the cross-correlation between variables).

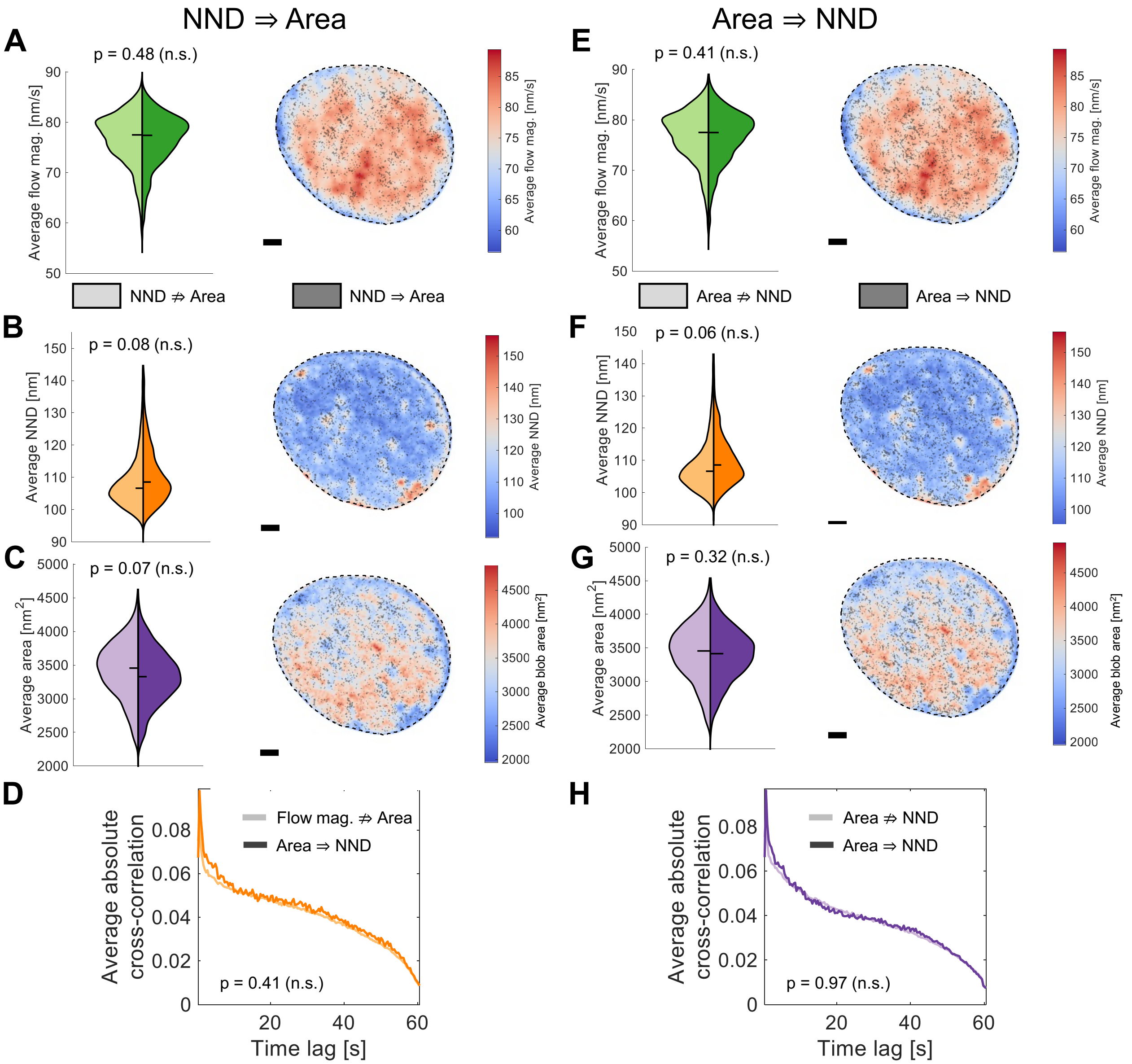


**Supplementary Figure 2: Additional parameter comparisons for the two cases that the NND causes the blob area (left) and vice versa (right).** Related to and in the same style of **Error! Reference source not found.** of the main text.

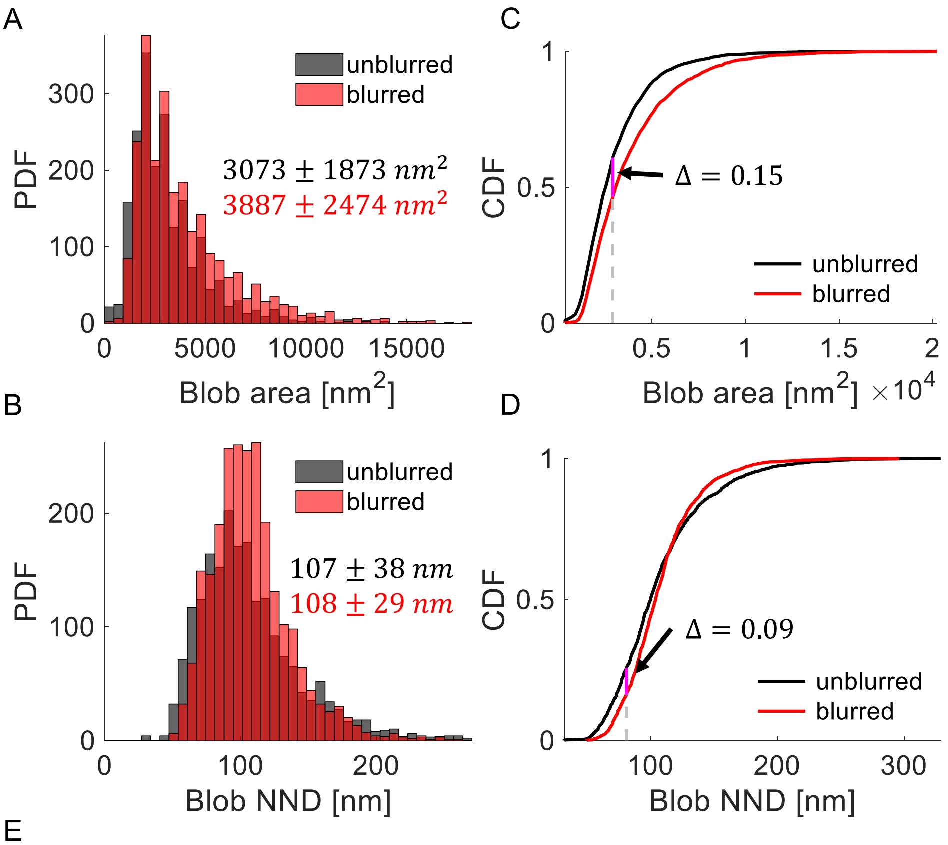


E

$$\frac{\text{Area}_{\text{blurred}} - \text{Area}_{\text{unblurred}}}{\mu_{\text{Area}}} = 0.23 \qquad \frac{\text{Area}_{\text{blurred}} - \text{Area}_{\text{unblurred}}}{\sigma_{\text{Area}}} = 0.37$$

$$\frac{\text{NND}_{\text{blurred}} - \text{NND}_{\text{unblurred}}}{\mu_{\text{NND}}} = 0.006 \qquad \frac{\text{NND}_{\text{blurred}} - \text{NND}_{\text{unblurred}}}{\sigma_{\text{NND}}} = 0.02$$

**Supplementary Figure 3: The blob NND is more robust against merging in regions with a density of blobs than the blob area.** Blobs on simulated images (as in [16]) were segmented on the original simulated frame and a blurred version of it. The blobs in the blurred and unblurred frames were characterized by **A)** their area and **B)** their NND. The empirical cumulative distribution function (CDF) was computed **C)** for the area and **D)** the NND and the maximum difference between the CDFs of the blobs identified in the blurred and unblurred images is indicated. A lower **Δ** indicates that the distributions in A-B) are more similar. **E)** Additional useful quantities were computed to examine how blob area and NND react to a blob merging. Shows are the scaled mean differences of mean area and NND in the blurred and unblurred case, respectively. The scaling is over the overall mean area/NND on the left and over the overall standard deviation over the area/NND distribution on the right. The difference in NND between blurred and unblurred versions of the frames are consistently smaller than for the area, indicating that the NND is more robust against resolution-mediated merging of blobs than the area.
